# SARS-CoV2 infection in farmed mink, Netherlands, April 2020

**DOI:** 10.1101/2020.05.18.101493

**Authors:** Nadia Oreshkova, Robert-Jan Molenaar, Sandra Vreman, Frank Harders, Bas B. Oude Munnink, Renate Hakze, Nora Gerhards, Paulien Tolsma, Ruth Bouwstra, Reina Sikkema, Mirriam Tacken, Myrna M.T. de Rooij, Eefke Weesendorp, Marc Engelsma, Christianne Bruschke, Lidwien A.M. Smit, Marion Koopmans, Wim H.M. van der Poel, Arjan Stegeman

## Abstract

In April 2020, respiratory disease and increased mortality were observed in farmed mink on two farms in the Netherlands. In both farms, at least one worker had been found positive for SARS-CoV-2. Necropsies of the mink revealed interstitial pneumonia, and organ and swab samples tested positive for SARS-CoV-2 RNA by qPCR. Variations in viral genomes point at between-mink transmission on the farms and lack of infection link between the farms. Inhalable dust in the mink houses contained viral RNA, indicating possible exposure of workers.

## Introduction

Currently, humanity is facing a pandemic of a new coronavirus, SARS-CoV-2 (Severe acute respiratory syndrome-coronavirus-2). The virus is spreading efficiently among people, causing predominantly respiratory disease with varying degree of severity. The virus was also shown to infect a number of animal species under experimental conditions. Rhesus and cynomolgus macaques, ferrets, cats and golden Syrian hamsters supported viral replication in respiratory tract [1-9] and some of those species (rhesus macaques, juvenile cats and hamsters) displayed a mild to moderate clinical disease. Besides the experimental infections, occasional spill-over from humans to domestic or captive animals has been reported. In a few isolated cases cats and dogs owned by infected individuals tested positive for SARS-CoV-2 RNA [10, 11], and occasionally, cats also displayed clinical disease. Recently, several tigers in the Bronx zoo with respiratory symptoms were confirmed positive for SARS-CoV-2. In all cases a direct correlation with infected humans was established, or at least other sources of infection were excluded [10].

Here, we report SARS-CoV-2 infection of mink on two farms in the Netherlands, and describe the associated clinical signs, pathological and virological findings. Sequence analysis of mink-derived viruses imply the role of SARS-CoV-2-positive humans as a probable source of the initial infection, point at transmission between mink and reveal possible exposure of workers to virus excreted by mink in the environment.

## Disease history and clinical observations

On 19 and 20 of April 2020, signs of respiratory disease were reported on two mink farms (NB1 with two closely situated locations: NB1a and NB1b, and NB2) in the Netherlands, province Noord Brabant. The respiratory signs were mostly limited to watery nasal discharge, but some animals showed severe respiratory distress. Overall mortality between date of reporting and 30 of April was 2.4% at NB1 and 1.2% at NB2, while approximately 0.6% would have been expected. Affected animals were spread throughout the farms. At this time of the year, the mink populations consist mainly of pregnant females. In the few litters that were already present, no increase in pup mortality was noticed.

Lungs from three recently died animals per farm were collected and submitted for qPCR analysis on 21 (NB1) and 25 (NB2) of April. One sample per farm was also sequenced (index samples). In the following week, thirty-six recently dead animals were collected (18 per farm) and necropsied. A throat and rectal swab were taken from each animal for qPCR analysis.

## Pathological analysis

### Macroscopic findings

The necropsies revealed that 16/18 animals from NB1 and 12/18 from NB2 had diffusely dark to mottled red, wet lung lobes that did not collapse when opening the thoracic cavity, indicating interstitial pneumonia (Table 1 and Figure 1A). Other investigated organs displayed no significant macroscopic changes. Mink without the described lung findings had macroscopic changes consistent with either chronic Aleutian disease, septicaemia, or dystocia. From 7 animals with clear macroscopic lung changes, organs were harvested for histopathological and virological investigation.

**Table 1.** Gross pathology/ cause of death overview of the necropsied mink

| Farm NB1 |  |  |  |  | Farm NB2 |  |  |  |
| --- | --- | --- | --- | --- | --- | --- | --- | --- |
| Animal # | NB1a or NB1b | Date of death | Date of necropsy | Cause of death | Animal # | Date of death | Date of necropsy | Cause of death |
| 1 | NB1a | 28/4/20 | 28/4/20 | interstitial pneumonia | 1 | 27/4/20 | 27/4/20 | sepsis and lung edema with congestion |
| 2 | NB1a | 28/4/20 | 28/4/20 | interstitial pneumonia* | 2 | 27/4/20 | 27/4/20 | interstitial pneumonia* |
| 3 | NB1a | 28/4/20 | 28/4/20 | interstitial pneumonia | 3 | 27/4/20 | 27/4/20 | Aleutian disease |
| 4 | NB1a | 28/4/20 | 28/4/20 | interstitial pneumonia | 4 | 27/4/20 | 27/4/20 | Aleutian disease |
| 5 | NB1a | 28/4/20 | 28/4/20 | interstitial pneumonia | 5 | 27/4/20 | 27/4/20 | sepsis |
| 6 | NB1a | 28/4/20 | 28/4/20 | interstitial pneumonia | 6 | 27/4/20 | 27/4/20 | dystocia |
| 7 | NB1a | 28/4/20 | 28/4/20 | interstitial pneumonia | 7 | 27/4/20 | 27/4/20 | interstitial pneumonia* |
| 8 | NB1a | 28/4/20 | 28/4/20 | interstitial pneumonia | 8 | 27/4/20 | 27/4/20 | interstitial pneumonia* |
| 9 | NB1a | 28/4/20 | 28/4/20 | Aleutian disease | 9 | 26/4/20 | 27/4/20 | interstitial pneumonia |
| 10 | NB1a | 28/4/20 | 28/4/20 | interstitial pneumonia | 10 | 26/4/20 | 27/4/20 | interstitial pneumonia |
| 11 | NB1a | 28/4/20 | 28/4/20 | interstitial pneumonia | 11 | 26/4/20 | 27/4/20 | interstitial pneumonia |
| 12 | NB1a | 28/4/20 | 28/4/20 | interstitial pneumonia | 12 | 26/4/20 | 27/4/20 | interstitial pneumonia |
| **13** | NB1a | 28/4/20 | 28/4/20 | interstitial pneumonia* | **13** | 26/4/20 | 27/4/20 | interstitial pneumonia |
| **14** | NB1b | 28/4/20 | 28/4/20 | interstitial pneumonia* | **14** | 26/4/20 | 27/4/20 | interstitial pneumonia |
| **15** | NB1b | 28/4/20 | 28/4/20 | interstitial pneumonia | **15** | 26/4/20 | 27/4/20 | interstitial pneumonia |
| **16** | NB1b | 28/4/20 | 28/4/20 | interstitial pneumonia* | **16** | 26/4/20 | 27/4/20 | interstitial pneumonia |
| **17** | NB1b | 28/4/20 | 28/4/20 | interstitial pneumonia | **17** | 26/4/20 | 27/4/20 | interstitial pneumonia |
| **18** | NB1b | 28/4/20 | 28/4/20 | interstitial pneumonia | **18** | 26/4/20 | 27/4/20 | sepsis |

\*Organs from those animals were collected for SARS-CoV-2 qPCR

**Figure 1.**
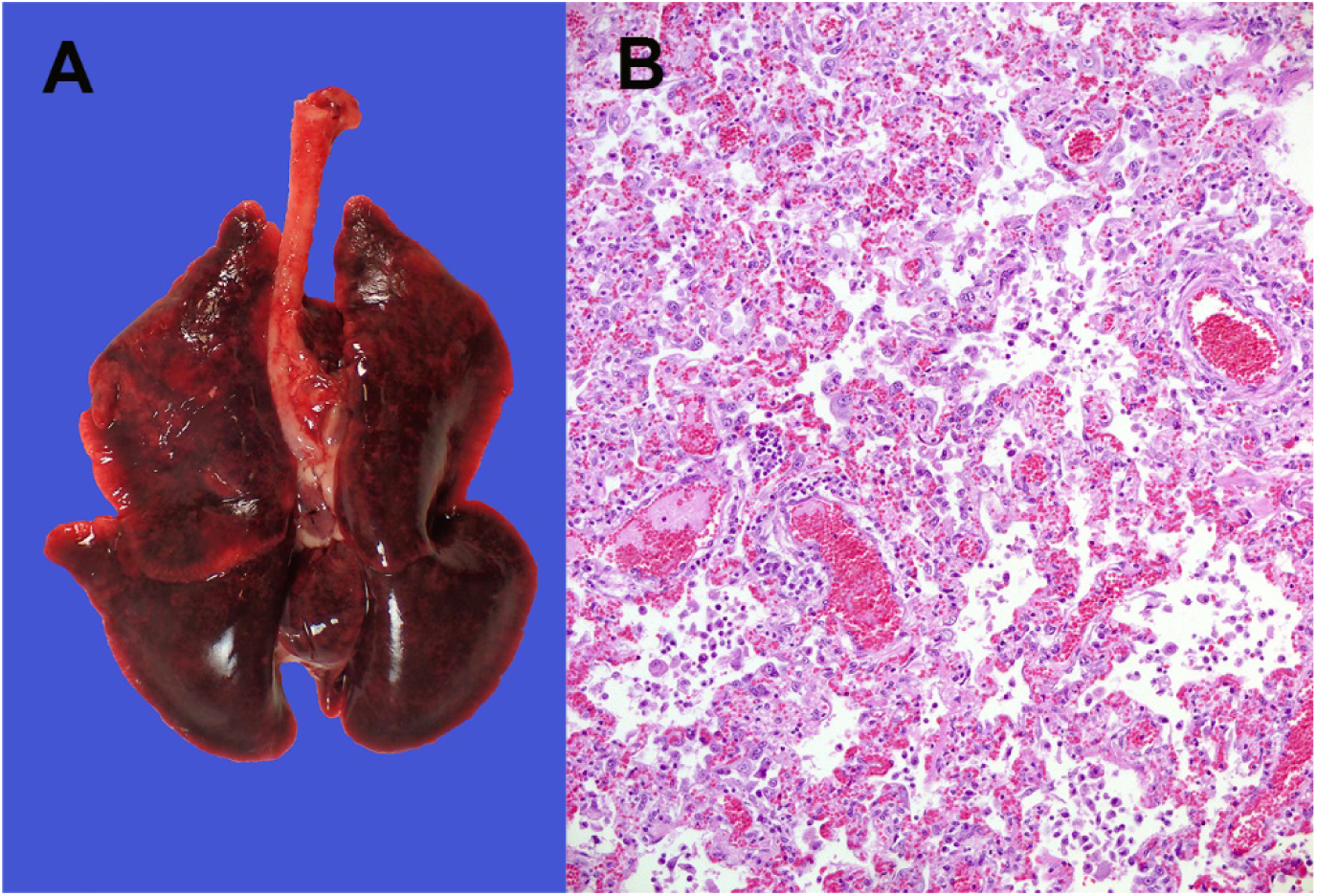
Lung from a necropsied mink. A) Representative macroscopic image of an affected lung and B) Representative microscopic image (objective 20x) of a section of the lung, fixed in 10% formalin and stained with haematoxylin and eosin (HE), showing interstitial pneumonia.

### Histological findings

A severe diffuse interstitial pneumonia with hyperaemia, alveolar damage and loss of air containing alveolar lumina was detected in all the 7 harvested lungs (Figure 1B). Bacterial cultures from the organs of the 7 animals were negative.

## Virus detection and sequencing

Presence of viral RNA was determined by qPCR against the E gene (Table 2) [12]. Viral RNA was detected in the conchae, lung, throat swab and rectal swab of all seven mink from which organs were collected. In addition, viral RNA was detected in the liver of one, and in the intestines of three animals. Spleens of all eight animals were negative for viral RNA (Table 2). In the swabs collected from all 36 necropsied animals, viral RNA was detected in all throat swabs and 34/36 of the rectal swabs. The Ct values varied, but on average were lower in the throat swabs, as compared to the rectal swabs (average Ct 21.7 and 31.2 respectively), indicating higher viral loads in the throat swabs.

**Table 2.** Virus titers, determined by qPCR in organs and swabs of necropsied mink. Titers were calculated based on a calibration curve of a virus stock with a known infectious virus titer and are expressed as log_10_ TCID50/gram tissue (organ material) or per ml of swab material (swabs were always submerged in 2 ml of cell culture medium).

|  | **Animal #** | **conchae** | **Lung** | **spleen** | **liver** | **intestines** | **throat swab** | **rectal swab** |
| --- | --- | --- | --- | --- | --- | --- | --- | --- |
| **Farm NB2** | **2** | 8,19 | 5,77 | not detected | not detected | not detected | 8,03 | 2,58 |
|  | **8** | 8,55 | 5,55 | not detected | 3,45 | not detected | 7,30 | 3,84 |
|  | **7** | 8,46 | 5,98 | not detected | not detected | not detected | 6,69 | 5,42 |
| **Farm NB1** | **2** | 8,25 | 4,54 | not detected | not detected | 4,22 | 6,87 | 3,30 |
|  | **13** | 9,16 | 5,17 | not detected | not detected | 3,56 | 6,81 | 3,01 |
|  | **14** | 8,08 | 3,83 | not detected | not detected | not detected | 7,04 | 3,95 |
|  | **16** | 7,08 | 3,90 | not detected | not detected | 4,97 | 6,47 | 4,47 |

The viral sequences of the index samples and from additional four and five animals from NB1 and NB2 respectively, were determined by NGS and deposited in GenBank (MT396266 and MT457390 – MT457399). Phylogenetic analysis of the sequences suggests two separate introductions into the two farms (Figure 2). The index sequences show 9 (NB1) and 15 (NB2) nucleotide substitutions over the complete genome in comparison with Wuhan-Hu-1 (NC_045512.2, EPI_ISL_402125). The two index sequences diverge at 22 nucleotide positions, but the sequences from each farm cluster together. A deletion of 3 nucleotides at positions 1605-7 of all sequences from NB2 resulted in a deletion of an aspartic acid from Orf1ab (Asp447-).

**Figure 2.**
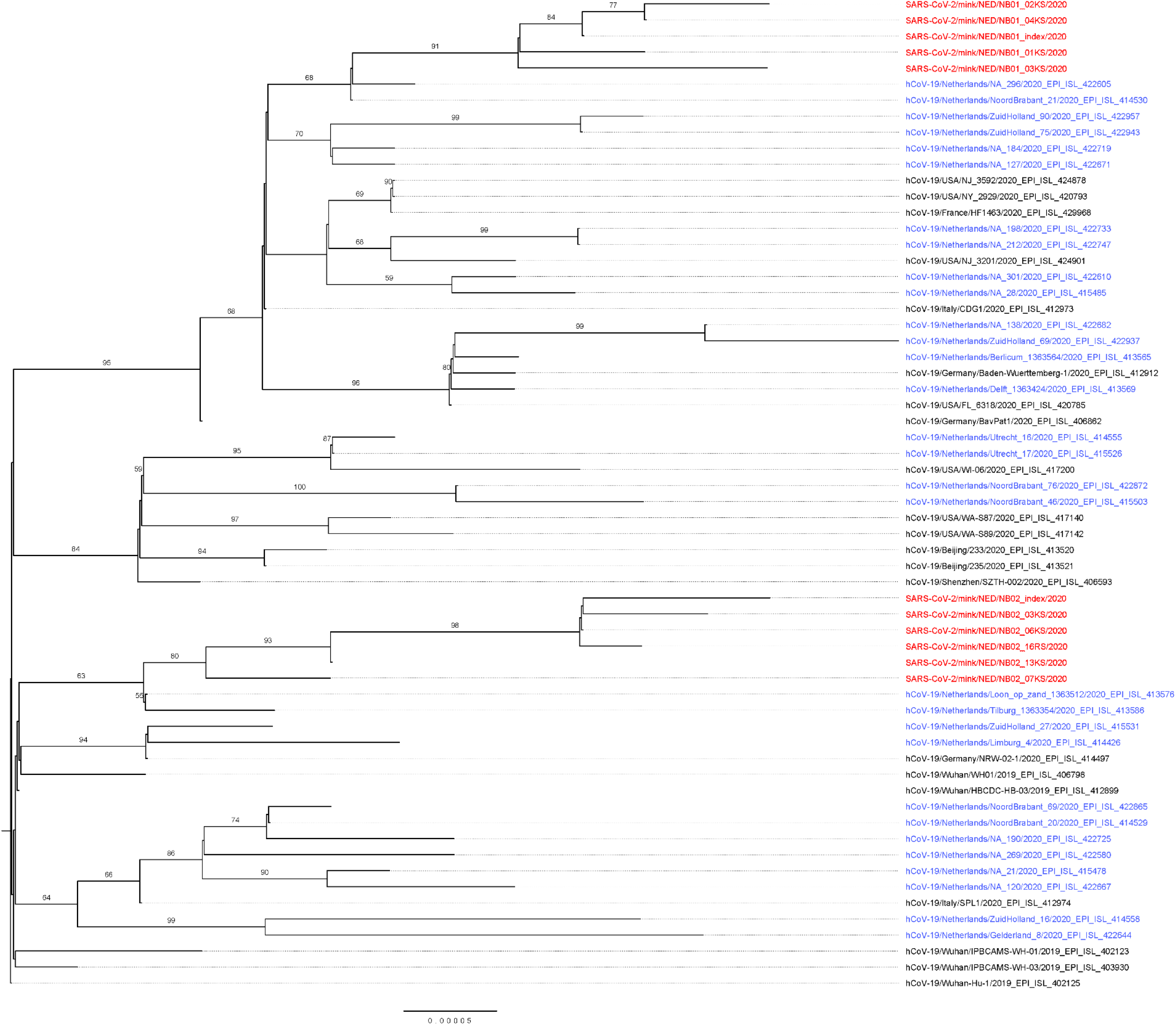
Maximum likelihood phylogenetic tree based on the full length SARS-CoV-2 sequences form mink (in red) and selected sequences, dutch (in blue) and international, from the GISAID EpiCoV Database (https://www.gisaid.org/; details and acknowledgements in supplementary table 1) based on proposed lineages [13]. The collected sequences were aligned using MAFFT v7.427 [14] and the evolutionary history was inferred by RAxML version 8.2.12 [15] utilizing the Maximum Likelihood method based on the General Time Reversible model with a gamma-distributed variation of rates and 1000 bootstrap replicates. The tree is rooted at Wuhan-Hu-1. Bootstrap support values above 50 are indicated at the corresponding branch.

## Covid-19 history in farm workers

Farm owners and their families were interviewed by the public health service for possible history of disease. One person on farm NB1 had disease symptoms since beginning of April, but was not investigated for SARS-CoV-2 infection. At NB2, one worker had been diagnosed with SARS-CoV-2 infection and hospitalized on 31^st^ of March 31. A clinical sample was retrieved, but the viral load was too low for sequencing analysis. The follow-up investigation is ongoing and will be reported elsewhere.

## Environmental sampling

Inhalable dust in the air was collected at three locations in each of the mink houses between 28^th^ of April and 2^nd^ of May by active stationary sampling during 5-6 hours, using Gilian GilAir 5 pumps at 3.5 L/min, GSP sampling heads and Teflon filters. Viral RNA was detected in 2/3 samples of NB1a (Ct 35.95 and 38.18), 1/3 of NB1b (Ct 35.03) and 1/3 of NB2 (Ct 35.14).

## Discussion

Here we present the first report of infection of two mink farms with SARS-CoV-2. In both farms, Covid-19 like symptoms were present in individuals working on the farms before signs in mink were seen, and infection was confirmed in one hospitalized person. The sequences obtained from the two farms are closely related to known human sequences, and the distance between the sequence clusters from the two farms suggests separate introductions. The most likely explanation of the widespread infection in the mink farms is introduction of the virus by humans and subsequent transmission amongst the minks. In the mink farms, the animals are caged separately with non-permeable partition between cages, precluding direct contact as a mode of transmission. Indirect transmission between minks could either be through fomites (e.g. by feed or bedding material provided by humans), by infectious droplets generated by the infected animals, or by (fecally-) contaminated dust from the bedding. Detection of viral RNA in the airborne inhalable dust in the mink farms clearly suggests dust and/or droplets as means of transmission between the minks and occupational risk of exposure for the workers on the farms.

In conclusion, this report shows the first SARS-CoV-2 virus outbreak in farmed mink. We demonstrate that mink are susceptible to infection with SARS-CoV-2 virus, may develop respiratory disease with typical pathological findings of viral pneumonia, and can transmit the virus amongst each other. Moreover, humans working in the mink houses are exposed to virus, indicating the need for infection prevention among workers and other biosecurity measures to prevent onward spread of the virus from the affected mink farms.

## Notes

### Competing Interest Statement

The authors have declared no competing interest.

