## Supplemental table 1 for "SARS-CoV2 infection in farmed mink, Netherlands, April 2020"

**We gratefully acknowledge the authors, originating and submitting laboratories of the sequences from GISAID's EpiCoV™ Database on which this research is based. The list is detailed below.**

**All submitters of data may be contacted directly via www.gisaid.org**

| Accession ID | Virus name | Location | Collection date | Originating lab | Submitting lab | Authors |
|---|---|---|---|---|---|---|
| EPI_ISL_402123 | hCoV-19/Wuhan/IPBCAMS-WH-01/2019 | Asia / China / Hubei / Wuhan | 2019-12-24 | Institute of Pathogen Biology, Chinese Academy of Medical Sciences & Peking Union Medical College | Institute of Pathogen Biology, Chinese Academy of Medical Sciences & Peking Union Medical College | Lili Ren, Jianwei Wang, Qi Jin, Zichun Xiang, Zhiqiang Wu, Chao Wu, Yiwei Liu |
| EPI_ISL_402125 | hCoV-19/Wuhan-Hu-1/2019 | Asia / China | 2019-12-31 | unknown | National Institute for Communicable Disease Control and Prevention (ICDC) Chinese Center for Disease Control and Prevention (China CDC) | Zhang,Y.-Z., Wu,F., Chen,Y.-M., Pei,Y.-Y., Xu,L., Wang,W., Zhao,S., Yu,B., Hu,Y., Tao,Z.-W., Song,Z.-G., Tian,J.-H., Zhang,Y.-L., Liu,Y., Zheng,J.-J., Dai,F.-H., Wang,Q.-M., She,J.-L. and Zhu,T.-Y. |
| EPI_ISL_403930 | hCoV-19/Wuhan/IPBCAMS-WH-03/2019 | Asia / China / Hubei / Wuhan | 2019-12-30 | Institute of Pathogen Biology, Chinese Academy of Medical Sciences & Peking Union Medical College | Institute of Pathogen Biology, Chinese Academy of Medical Sciences & Peking Union Medical College | Lili Ren, Jianwei Wang, Qi Jin, Zichun Xiang, Zhiqiang Wu, Chao Wu, Yiwei Liu |
| EPI_ISL_406593 | hCoV-19/Shenzhen/SZTH-002/2020 | Asia / China / Guandong / Shenzhen | 2020-01-13 | Shenzhen Key Laboratory of Pathogen and Immunity, National Clinical Research Center for Infectious Disease, Shenzhen Third People's Hospital | Shenzhen Key Laboratory of Pathogen and Immunity, National Clinical Research Center for Infectious Disease, Shenzhen Third People's Hospital | Yang Yang, Chenguang Shen, Li Xing, Zhixiang Xu, Haixia Zheng, Yingxia Liu |
| EPI_ISL_406798 | hCoV-19/Wuhan/WH01/2019 | Asia / China / Hubei / Wuhan | 2019-12-26 | General Hospital of Central Theater Command of People's Liberation Army of China | BGI & Institute of Microbiology, Chinese Academy of Sciences & Shandong First Medical University & Shandong Academy of Medical Sciences & General Hospital of Central Theater Command of People's Liberation Army of China | Weijun Chen, Yuhai Bi, Weifeng Shi and Zhenhong Hu |
| EPI_ISL_406862 | hCoV-19/Germany/BavPat1/2020 | Europe / Germany / Bavaria / Munich | 2020-01-28 | Charité Universitätsmedizin Berlin, Institute of Virology; Institut für Mikrobiologie der Bundeswehr, Munich | Charité Universitätsmedizin Berlin, Institute of Virology | Victor M Corman, Julia Schneider, Talitha Veith, Barbara Mühlemann, Markus Antwerpen, Christian Drosten, Roman Wölfel |
| EPI_ISL_412899 | hCoV-19/Wuhan/HBCDC-HB-03/2019 | Asia / China / Hubei / Wuhan | 2019-12-30 | Wuhan Jinyintan Hospital | Hubei Provincial Center for Disease Control and Prevention | Bin Fang, Xiang Li, Xiao Yu, Linlin Liu, Bo Yang, Faxian Zhan, Guojun Ye, Xixiang Huo, Junqiang Xu, Bo Yu, Kun Cai, Jing Li, Yongzhong Jiang. |
| EPI_ISL_412912 | hCoV-19/Germany/Baden-Wuerttemberg-1/2020 | Europe / Germany / Baden-Wuerttemberg | 2020-02-25 | State Health Office Baden-Wuerttemberg | Charité Universitätsmedizin Berlin, Institute of Virology | Victor M Corman, Julia Schneider, Barbara Mühlemann, Talitha Veith, Jörn Beheim-Schwarzbach, Terry Jones, Rainer Oehme, Silke Fischer, Christian Drosten |
| EPI_ISL_412973 | hCoV-19/Italy/CDG1/2020 | Europe / Italy / Lombardy | 2020-02-20 | Department of Infectious Diseases, Istituto Superiore di Sanità, Roma , Italy | Virology Laboratory, Scientific Department, Army Medical Center | Paola Stefanelli, Stefano Fiore, Antonella Marchi, Eleonora Benedetti, Concetta Fabiani, Giovanni Faggioni, Antonella Fortunato, Riccardo De Santis, Silvia Fillo, Anna Anselmo, Andrea Ciammaruconi, Stefano Palomba, Florigio Lista |
| EPI_ISL_412974 | hCoV-19/Italy/SPL1/2020 | Europe / Italy / Rome | 2020-01-29 | Department of Infectious Diseases, Istituto Superiore di Sanità, Rome, Italy | Virology Laboratory, Scientific Department, Army Medical Center | Paola Stefanelli, Stefano Fiore, Antonella Marchi, Eleonora Benedetti, Concetta Fabiani, Giovanni Faggioni, Antonella Fortunato, Silvia Fillo, Riccardo De Santis, Andrea Ciammaruconi, Giancarlo Petralito, Filippo Molinari, Florigio Lista |
| EPI_ISL_413520 | hCoV-19/Beijing/233/2020 | Asia / China / Beijing | 2020-01-28 | unknown | Infectious Disease Control Center | Li,J., Li,L., Li,Z., Qiu,S., Song,H., Li,P. and Li,P. |
| EPI_ISL_413521 | hCoV-19/Beijing/235/2020 | Asia / China / Beijing | 2020-01-28 | unknown | Infectious Disease Control Center | Li,J., Li,L., Li,Z., Qiu,S., Song,H., Li,P. and Li,P. |
| EPI_ISL_413565 | hCoV-19/Netherlands/Berlicum_1363564/2020 | Europe / Netherlands / Berlicum | 2020-02-24 | Foundation Pamm | Erasmus Medical Center | David Nieuwenhuijse, Bas Oude Munnink, Reina Sikkema, Claudia Schapendonk, Irina Chestakova, Anne van der Linden, Mark Pronk, Pascal Lexmond, Corien Swaan, Manon Haverkate, Madelief Mollers, Mart Stein, Sandra Kengne Kamga Mobou, Jeroen van Kampen, Jolanda Voermans, Aura Timen, Corine GeurtsvanKessel, Annemiek van der Eijk, Richard Molenkamp, Marion Koopmans, on behalf of the Dutch national COVID-19 response team. |
| EPI_ISL_413569 | hCoV-19/Netherlands/Delft_1363424/2020 | Europe / Netherlands / Delft | 2020-02-28 | RIVM | Erasmus Medical Center | David Nieuwenhuijse, Bas Oude Munnink, Reina Sikkema, Claudia Schapendonk, Irina Chestakova, Anne van der Linden, Mark Pronk, Pascal Lexmond, Corien Swaan, Manon Haverkate, Madelief Mollers, Mart Stein, Sandra Kengne Kamga Mobou, Jeroen van Kampen, Jolanda Voermans, Aura Timen, Corine GeurtsvanKessel, Annemiek van der Eijk, Richard Molenkamp, Marion Koopmans, on behalf of the Dutch national COVID-19 response team. |
| EPI_ISL_413576 | hCoV-19/Netherlands/Loon_op_zand_1363512/2020 | Europe / Netherlands / Loon op zand | 2020-02-29 | RIVM | Erasmus Medical Center | David Nieuwenhuijse, Bas Oude Munnink, Reina Sikkema, Claudia Schapendonk, Irina Chestakova, Anne van der Linden, Mark Pronk, Pascal Lexmond, Corien Swaan, Manon Haverkate, Madelief Mollers, Mart Stein, Sandra Kengne Kamga Mobou, Jeroen van Kampen, Jolanda Voermans, Aura Timen, Corine GeurtsvanKessel, Annemiek van der Eijk, Richard Molenkamp, Marion Koopmans, on behalf of the Dutch national COVID-19 response team. |
| EPI_ISL_413586 | hCoV-19/Netherlands/Tilburg_1363354/2020 | Europe / Netherlands / Tilburg | 2020-02-27 | Foundation Elisabeth-Tweesteden Ziekenhuis | Erasmus Medical Center | David Nieuwenhuijse, Bas Oude Munnink, Reina Sikkema, Claudia Schapendonk, Irina Chestakova, Anne van der Linden, Mark Pronk, Pascal Lexmond, Corien Swaan, Manon Haverkate, Madelief Mollers, Mart Stein, Sandra Kengne Kamga Mobou, Jeroen van Kampen, Jolanda Voermans, Aura Timen, Corine GeurtsvanKessel, Annemiek van der Eijk, Richard Molenkamp, Marion Koopmans, on behalf of the Dutch national COVID-19 response team. |

| | | | | | | |
|---|---|---|---|---|---|---|
| EPI_ISL_414426 | hCoV-19/Netherlands/Limburg_4/2020 | Europe / Netherlands / Limburg | 2020-03-03 | Dutch COVID-19 response team | Erasmus Medical Center | David Nieuwenhuijse, Bas Oude Munnink, Reina Sikkema, Claudia Schapendonk, Irina Chestakova, Anne van der Linden, Mark Pronk, Pascal Lexmond, Corien Swaan, Manon Haverkate, Madelief Mollers, Mart Stein, Sandra Kengne Kamga Mobou, Jeroen van Kampen, Jolanda Voermans, Aura Timen, Corine GeurtsvanKessel, Annemiek van der Eijk, Richard Molenkamp, Marion Koopmans, on behalf of the Dutch national COVID-19 response team. |
| EPI_ISL_414497 | hCoV-19/Germany/NRW-02-1/2020 | Europe / Germany / North Rhine Westphalia / Heinsberg District | 2020-02-25 | Center of Medical Microbiology, Virology, and Hospital Hygiene, University of Duesseldorf | Center of Medical Microbiology, Virology, and Hospital Hygiene, University of Duesseldorf | Ortwin Adams, Marcel Andree, Alexander Dilthey, Torsten Feldt, Sandra Hauka, Torsten Houwaart, Björn-Erik Jensen, Detlef Kindgen-Milles, Malte Kohns Vasconcelos, Klaus Pfeffer, Tina Senff, Daniel Strelow, Jörg Timm, Andreas Walker, Tobias Wienemann |
| EPI_ISL_414529 | hCoV-19/Netherlands/NoordBrabant_20/2020 | Europe / Netherlands / Noord Brabant | 2020-03-04 | Dutch COVID-19 response team | Erasmus Medical Center | David Nieuwenhuijse, Bas Oude Munnink, Reina Sikkema, Claudia Schapendonk, Irina Chestakova, Anne van der Linden, Mark Pronk, Pascal Lexmond, Corien Swaan, Manon Haverkate, Madelief Mollers, Mart Stein, Sandra Kengne Kamga Mobou, Jeroen van Kampen, Jolanda Voermans, Aura Timen, Corine GeurtsvanKessel, Annemiek van der Eijk, Richard Molenkamp, Marion Koopmans, on behalf of the Dutch national COVID-19 response team. |
| EPI_ISL_414530 | hCoV-19/Netherlands/NoordBrabant_21/2020 | Europe / Netherlands / Noord Brabant | 2020-03-04 | Dutch COVID-19 response team | Erasmus Medical Center | David Nieuwenhuijse, Bas Oude Munnink, Reina Sikkema, Claudia Schapendonk, Irina Chestakova, Anne van der Linden, Mark Pronk, Pascal Lexmond, Corien Swaan, Manon Haverkate, Madelief Mollers, Mart Stein, Sandra Kengne Kamga Mobou, Jeroen van Kampen, Jolanda Voermans, Aura Timen, Corine GeurtsvanKessel, Annemiek van der Eijk, Richard Molenkamp, Marion Koopmans, on behalf of the Dutch national COVID-19 response team. |
| EPI_ISL_414555 | hCoV-19/Netherlands/Utrecht_16/2020 | Europe / Netherlands / Utrecht | 2020-03-08 | Dutch COVID-19 response team | Erasmus Medical Center | David Nieuwenhuijse, Bas Oude Munnink, Reina Sikkema, Claudia Schapendonk, Irina Chestakova, Anne van der Linden, Mark Pronk, Pascal Lexmond, Corien Swaan, Manon Haverkate, Madelief Mollers, Mart Stein, Sandra Kengne Kamga Mobou, Jeroen van Kampen, Jolanda Voermans, Aura Timen, Corine GeurtsvanKessel, Annemiek van der Eijk, Richard Molenkamp, Marion Koopmans, on behalf of the Dutch national COVID-19 response team. |
| EPI_ISL_414558 | hCoV-19/Netherlands/ZuidHolland_16/2020 | Europe / Netherlands / Zuid Holland | 2020-03-06 | Dutch COVID-19 response team | Erasmus Medical Center | David Nieuwenhuijse, Bas Oude Munnink, Reina Sikkema, Claudia Schapendonk, Irina Chestakova, Anne van der Linden, Mark Pronk, Pascal Lexmond, Corien Swaan, Manon Haverkate, Madelief Mollers, Mart Stein, Sandra Kengne Kamga Mobou, Jeroen van Kampen, Jolanda Voermans, Aura Timen, Corine GeurtsvanKessel, Annemiek van der Eijk, Richard Molenkamp, Marion Koopmans, on behalf of the Dutch national COVID-19 response team. |
| EPI_ISL_415478 | hCoV-19/Netherlands/NA_21/2020 | Europe / Netherlands | 2020-03-08 | Dutch COVID-19 response team | Erasmus Medical Center | David Nieuwenhuijse, Bas Oude Munnink, Reina Sikkema, Claudia Schapendonk, Irina Chestakova, Anne van der Linden, Mark Pronk, Pascal Lexmond, Corien Swaan, Manon Haverkate, Madelief Mollers, Mart Stein, Sandra Kengne Kamga Mobou, Jeroen van Kampen, Jolanda Voermans, Aura Timen, Corine GeurtsvanKessel, Annemiek van der Eijk, Richard Molenkamp, Marion Koopmans, on behalf of the Dutch national COVID-19 response team. |
| EPI_ISL_415485 | hCoV-19/Netherlands/NA_28/2020 | Europe / Netherlands | 2020-03-12 | Dutch COVID-19 response team | Erasmus Medical Center | David Nieuwenhuijse, Bas Oude Munnink, Reina Sikkema, Claudia Schapendonk, Irina Chestakova, Anne van der Linden, Mark Pronk, Pascal Lexmond, Corien Swaan, Manon Haverkate, Madelief Mollers, Mart Stein, Sandra Kengne Kamga Mobou, Jeroen van Kampen, Jolanda Voermans, Aura Timen, Corine GeurtsvanKessel, Annemiek van der Eijk, Richard Molenkamp, Marion Koopmans, on behalf of the Dutch national COVID-19 response team. |
| EPI_ISL_415503 | hCoV-19/Netherlands/NoordBrabant_46/2020 | Europe / Netherlands / Noord Brabant | 2020-03-11 | Dutch COVID-19 response team | Erasmus Medical Center | David Nieuwenhuijse, Bas Oude Munnink, Reina Sikkema, Claudia Schapendonk, Irina Chestakova, Anne van der Linden, Mark Pronk, Pascal Lexmond, Corien Swaan, Manon Haverkate, Madelief Mollers, Mart Stein, Sandra Kengne Kamga Mobou, Jeroen van Kampen, Jolanda Voermans, Aura Timen, Corine GeurtsvanKessel, Annemiek van der Eijk, Richard Molenkamp, Marion Koopmans, on behalf of the Dutch national COVID-19 response team. |
| EPI_ISL_415526 | hCoV-19/Netherlands/Utrecht_17/2020 | Europe / Netherlands / Utrecht | 2020-03-10 | Dutch COVID-19 response team | Erasmus Medical Center | David Nieuwenhuijse, Bas Oude Munnink, Reina Sikkema, Claudia Schapendonk, Irina Chestakova, Anne van der Linden, Mark Pronk, Pascal Lexmond, Corien Swaan, Manon Haverkate, Madelief Mollers, Mart Stein, Sandra Kengne Kamga Mobou, Jeroen van Kampen, Jolanda Voermans, Aura Timen, Corine GeurtsvanKessel, Annemiek van der Eijk, Richard Molenkamp, Marion Koopmans, on behalf of the Dutch national COVID-19 response team. |

| | | | | | | |
|---|---|---|---|---|---|---|
| EPI_ISL_415531 | hCoV-19/Netherlands/ZuidHolland_27/2020 | Europe / Netherlands / Zuid Holland | 2020-03-09 | Dutch COVID-19 response team | Erasmus Medical Center | David Nieuwenhuijse, Bas Oude Munnink, Reina Sikkema, Claudia Schapendonk, Irina Chestakova, Anne van der Linden, Mark Pronk, Pascal Lexmond, Corien Swaan, Manon Haverkate, Madelief Mollers, Mart Stein, Sandra Kengne Kamga Mobou, Jeroen van Kampen, Jolanda Voermans, Aura Timen, Corine GeurtsvanKessel, Annemiek van der Eijk, Richard Molenkamp, Marion Koopmans, on behalf of the Dutch national COVID-19 response team. |
| EPI_ISL_417140 | hCoV-19/USA/WA-S87/2020 | North America / USA / Washington / King County | 2020-03-01 | Washington State Department of Health | Seattle Flu Study | Chu etl al |
| EPI_ISL_417142 | hCoV-19/USA/WA-S89/2020 | North America / USA / Washington / Umatilla County | 2020-02-29 | Washington State Department of Health | Seattle Flu Study | Chu etl al |
| EPI_ISL_417200 | hCoV-19/USA/WI-06/2020 | North America / USA / Wisconsin | 2020-03-21 | University of Wisconsin-Madison AIDS Vaccine Research Laboratories | University of Wisconsin-Madison AIDS Vaccine Research Laboratories | Katarina Braun and Gage Moreno |
| EPI_ISL_420785 | hCoV-19/USA/FL_6318/2020 | North America / USA / Florida | 2020-03-02 | FL Bureau of Health Laboratories Tampa | Pathogen Discovery, Respiratory Viruses Branch, Division of Viral Diseases, Centers for Disease Control and Prevention | Krista Queen, Yan Li, Ying Tao, Jing Zhang, Anne Uehara, Clinton R. Paden, Haibin Wang, Rachel Marine, Mary S. Keckler, Alison S. Laufer Halpin, Jasmine Padilla, Justin Lee, Christopher A. Elkins, Suxiang Tong |
| EPI_ISL_420793 | hCoV-19/USA/NY_2929/2020 | North America / USA / New York | 2020-03-02 | NYC Department of Health and Mental Hygiene | Pathogen Discovery, Respiratory Viruses Branch, Division of Viral Diseases, Centers for Disease Control and Prevention | Krista Queen, Yan Li, Ying Tao, Jing Zhang, Anne Uehara, Clinton R. Paden, Haibin Wang, Rachel Marine, Mary S. Keckler, Alison S. Laufer Halpin, Jasmine Padilla, Justin Lee, Christopher A. Elkins, Suxiang Tong |
| EPI_ISL_422580 | hCoV-19/Netherlands/NA_269/2020 | Europe / Netherlands | 2020-03-30 | Dutch COVID-19 response team | Erasmus Medical Center | Bas Oude Munnink, David Nieuwenhuijse, Reina Sikkema, Claudia Schapendonk, Irina Chestakova, Anne van der Linden, Theo Bestebroer, Stefan van Nieuwkoop, Mark Pronk, Pascal Lexmond, Corien Swaan, Manon Haverkate, Madelief Mollers, Mart Stein, Sandra Kengne Kamga Mobou, Jeroen van Kampen, Jolanda Voermans, Aura Timen, Corine GeurtsvanKessel, Annemiek van der Eijk, Richard Molenkamp, Marion Koopmans, on behalf of the Dutch national COVID-19 response team. |
| EPI_ISL_422605 | hCoV-19/Netherlands/NA_296/2020 | Europe / Netherlands | 2020-04-01 | Dutch COVID-19 response team | Erasmus Medical Center | Bas Oude Munnink, David Nieuwenhuijse, Reina Sikkema, Claudia Schapendonk, Irina Chestakova, Anne van der Linden, Theo Bestebroer, Stefan van Nieuwkoop, Mark Pronk, Pascal Lexmond, Corien Swaan, Manon Haverkate, Madelief Mollers, Mart Stein, Sandra Kengne Kamga Mobou, Jeroen van Kampen, Jolanda Voermans, Aura Timen, Corine GeurtsvanKessel, Annemiek van der Eijk, Richard Molenkamp, Marion Koopmans, on behalf of the Dutch national COVID-19 response team. |
| EPI_ISL_422610 | hCoV-19/Netherlands/NA_301/2020 | Europe / Netherlands | 2020-04-01 | Dutch COVID-19 response team | Erasmus Medical Center | Bas Oude Munnink, David Nieuwenhuijse, Reina Sikkema, Claudia Schapendonk, Irina Chestakova, Anne van der Linden, Theo Bestebroer, Stefan van Nieuwkoop, Mark Pronk, Pascal Lexmond, Corien Swaan, Manon Haverkate, Madelief Mollers, Mart Stein, Sandra Kengne Kamga Mobou, Jeroen van Kampen, Jolanda Voermans, Aura Timen, Corine GeurtsvanKessel, Annemiek van der Eijk, Richard Molenkamp, Marion Koopmans, on behalf of the Dutch national COVID-19 response team. |
| EPI_ISL_422644 | hCoV-19/Netherlands/Gelderland_8/2020 | Europe / Netherlands / Gelderland | 2020-03-13 | Dutch COVID-19 response team | Erasmus Medical Center | Bas Oude Munnink, David Nieuwenhuijse, Reina Sikkema, Claudia Schapendonk, Irina Chestakova, Anne van der Linden, Theo Bestebroer, Stefan van Nieuwkoop, Mark Pronk, Pascal Lexmond, Corien Swaan, Manon Haverkate, Madelief Mollers, Mart Stein, Sandra Kengne Kamga Mobou, Jeroen van Kampen, Jolanda Voermans, Aura Timen, Corine GeurtsvanKessel, Annemiek van der Eijk, Richard Molenkamp, Marion Koopmans, on behalf of the Dutch national COVID-19 response team. |
| EPI_ISL_422667 | hCoV-19/Netherlands/NA_120/2020 | Europe / Netherlands | 2020-03-16 | Dutch COVID-19 response team | Erasmus Medical Center | Bas Oude Munnink, David Nieuwenhuijse, Reina Sikkema, Claudia Schapendonk, Irina Chestakova, Anne van der Linden, Theo Bestebroer, Stefan van Nieuwkoop, Mark Pronk, Pascal Lexmond, Corien Swaan, Manon Haverkate, Madelief Mollers, Mart Stein, Sandra Kengne Kamga Mobou, Jeroen van Kampen, Jolanda Voermans, Aura Timen, Corine GeurtsvanKessel, Annemiek van der Eijk, Richard Molenkamp, Marion Koopmans, on behalf of the Dutch national COVID-19 response team. |
| EPI_ISL_422671 | hCoV-19/Netherlands/NA_127/2020 | Europe / Netherlands | 2020-03-17 | Dutch COVID-19 response team | Erasmus Medical Center | Bas Oude Munnink, David Nieuwenhuijse, Reina Sikkema, Claudia Schapendonk, Irina Chestakova, Anne van der Linden, Theo Bestebroer, Stefan van Nieuwkoop, Mark Pronk, Pascal Lexmond, Corien Swaan, Manon Haverkate, Madelief Mollers, Mart Stein, Sandra Kengne Kamga Mobou, Jeroen van Kampen, Jolanda Voermans, Aura Timen, Corine GeurtsvanKessel, Annemiek van der Eijk, Richard Molenkamp, Marion Koopmans, on behalf of the Dutch national COVID-19 response team. |
| EPI_ISL_422682 | hCoV-19/Netherlands/NA_138/2020 | Europe / Netherlands | 2020-03-19 | Dutch COVID-19 response team | Erasmus Medical Center | Bas Oude Munnink, David Nieuwenhuijse, Reina Sikkema, Claudia Schapendonk, Irina Chestakova, Anne van der Linden, Theo Bestebroer, Stefan van Nieuwkoop, Mark Pronk, Pascal Lexmond, Corien Swaan, Manon Haverkate, Madelief Mollers, Mart Stein, Sandra Kengne Kamga Mobou, Jeroen van Kampen, Jolanda Voermans, Aura Timen, Corine GeurtsvanKessel, Annemiek van der Eijk, Richard Molenkamp, Marion Koopmans, on behalf of the Dutch national COVID-19 response team. |

| | | | | | | |
|---|---|---|---|---|---|---|
| EPI_ISL_422719 | hCoV-19/Netherlands/NA_184/2020 | Europe / Netherlands | 2020-03-22 | Dutch COVID-19 response team | Erasmus Medical Center | Bas Oude Munnink, David Nieuwenhuijse, Reina Sikkema, Claudia Schapendonk, Irina Chestakova, Anne van der Linden, Theo Bestebroer, Stefan van Nieuwkoop, Mark Pronk, Pascal Lexmond, Corien Swaan, Manon Haverkate, Madelief Mollers, Mart Stein, Sandra Kengne Kamga Mobou, Jeroen van Kampen, Jolanda Voermans, Aura Timen, Corine GeurtsvanKessel, Annemiek van der Eijk, Richard Molenkamp, Marion Koopmans, on behalf of the Dutch national COVID-19 response team. |
| EPI_ISL_422725 | hCoV-19/Netherlands/NA_190/2020 | Europe / Netherlands | 2020-03-23 | Dutch COVID-19 response team | Erasmus Medical Center | Bas Oude Munnink, David Nieuwenhuijse, Reina Sikkema, Claudia Schapendonk, Irina Chestakova, Anne van der Linden, Theo Bestebroer, Stefan van Nieuwkoop, Mark Pronk, Pascal Lexmond, Corien Swaan, Manon Haverkate, Madelief Mollers, Mart Stein, Sandra Kengne Kamga Mobou, Jeroen van Kampen, Jolanda Voermans, Aura Timen, Corine GeurtsvanKessel, Annemiek van der Eijk, Richard Molenkamp, Marion Koopmans, on behalf of the Dutch national COVID-19 response team. |
| EPI_ISL_422733 | hCoV-19/Netherlands/NA_198/2020 | Europe / Netherlands | 2020-03-24 | Dutch COVID-19 response team | Erasmus Medical Center | Bas Oude Munnink, David Nieuwenhuijse, Reina Sikkema, Claudia Schapendonk, Irina Chestakova, Anne van der Linden, Theo Bestebroer, Stefan van Nieuwkoop, Mark Pronk, Pascal Lexmond, Corien Swaan, Manon Haverkate, Madelief Mollers, Mart Stein, Sandra Kengne Kamga Mobou, Jeroen van Kampen, Jolanda Voermans, Aura Timen, Corine GeurtsvanKessel, Annemiek van der Eijk, Richard Molenkamp, Marion Koopmans, on behalf of the Dutch national COVID-19 response team. |
| EPI_ISL_422747 | hCoV-19/Netherlands/NA_212/2020 | Europe / Netherlands | 2020-03-24 | Dutch COVID-19 response team | Erasmus Medical Center | Bas Oude Munnink, David Nieuwenhuijse, Reina Sikkema, Claudia Schapendonk, Irina Chestakova, Anne van der Linden, Theo Bestebroer, Stefan van Nieuwkoop, Mark Pronk, Pascal Lexmond, Corien Swaan, Manon Haverkate, Madelief Mollers, Mart Stein, Sandra Kengne Kamga Mobou, Jeroen van Kampen, Jolanda Voermans, Aura Timen, Corine GeurtsvanKessel, Annemiek van der Eijk, Richard Molenkamp, Marion Koopmans, on behalf of the Dutch national COVID-19 response team. |
| EPI_ISL_422865 | hCoV-19/Netherlands/NoordBrabant_69/2020 | Europe / Netherlands / Noord Brabant | 2020-03-02 | Dutch COVID-19 response team | Erasmus Medical Center | Bas Oude Munnink, David Nieuwenhuijse, Reina Sikkema, Claudia Schapendonk, Irina Chestakova, Anne van der Linden, Theo Bestebroer, Stefan van Nieuwkoop, Mark Pronk, Pascal Lexmond, Corien Swaan, Manon Haverkate, Madelief Mollers, Mart Stein, Sandra Kengne Kamga Mobou, Jeroen van Kampen, Jolanda Voermans, Aura Timen, Corine GeurtsvanKessel, Annemiek van der Eijk, Richard Molenkamp, Marion Koopmans, on behalf of the Dutch national COVID-19 response team. |
| EPI_ISL_422872 | hCoV-19/Netherlands/NoordBrabant_76/2020 | Europe / Netherlands / Noord Brabant | 2020-03-09 | Dutch COVID-19 response team | Erasmus Medical Center | Bas Oude Munnink, David Nieuwenhuijse, Reina Sikkema, Claudia Schapendonk, Irina Chestakova, Anne van der Linden, Theo Bestebroer, Stefan van Nieuwkoop, Mark Pronk, Pascal Lexmond, Corien Swaan, Manon Haverkate, Madelief Mollers, Mart Stein, Sandra Kengne Kamga Mobou, Jeroen van Kampen, Jolanda Voermans, Aura Timen, Corine GeurtsvanKessel, Annemiek van der Eijk, Richard Molenkamp, Marion Koopmans, on behalf of the Dutch national COVID-19 response team. |
| EPI_ISL_422937 | hCoV-19/Netherlands/ZuidHolland_69/2020 | Europe / Netherlands / Zuid Holland | 2020-03-24 | Dutch COVID-19 response team | Erasmus Medical Center | Bas Oude Munnink, David Nieuwenhuijse, Reina Sikkema, Claudia Schapendonk, Irina Chestakova, Anne van der Linden, Theo Bestebroer, Stefan van Nieuwkoop, Mark Pronk, Pascal Lexmond, Corien Swaan, Manon Haverkate, Madelief Mollers, Mart Stein, Sandra Kengne Kamga Mobou, Jeroen van Kampen, Jolanda Voermans, Aura Timen, Corine GeurtsvanKessel, Annemiek van der Eijk, Richard Molenkamp, Marion Koopmans, on behalf of the Dutch national COVID-19 response team. |
| EPI_ISL_422943 | hCoV-19/Netherlands/ZuidHolland_75/2020 | Europe / Netherlands / Zuid Holland | 2020-03-26 | Dutch COVID-19 response team | Erasmus Medical Center | Bas Oude Munnink, David Nieuwenhuijse, Reina Sikkema, Claudia Schapendonk, Irina Chestakova, Anne van der Linden, Theo Bestebroer, Stefan van Nieuwkoop, Mark Pronk, Pascal Lexmond, Corien Swaan, Manon Haverkate, Madelief Mollers, Mart Stein, Sandra Kengne Kamga Mobou, Jeroen van Kampen, Jolanda Voermans, Aura Timen, Corine GeurtsvanKessel, Annemiek van der Eijk, Richard Molenkamp, Marion Koopmans, on behalf of the Dutch national COVID-19 response team. |
| EPI_ISL_422957 | hCoV-19/Netherlands/ZuidHolland_90/2020 | Europe / Netherlands / Zuid Holland | 2020-03-29 | Dutch COVID-19 response team | Erasmus Medical Center | Bas Oude Munnink, David Nieuwenhuijse, Reina Sikkema, Claudia Schapendonk, Irina Chestakova, Anne van der Linden, Theo Bestebroer, Stefan van Nieuwkoop, Mark Pronk, Pascal Lexmond, Corien Swaan, Manon Haverkate, Madelief Mollers, Mart Stein, Sandra Kengne Kamga Mobou, Jeroen van Kampen, Jolanda Voermans, Aura Timen, Corine GeurtsvanKessel, Annemiek van der Eijk, Richard Molenkamp, Marion Koopmans, on behalf of the Dutch national COVID-19 response team. |
| EPI_ISL_424878 | hCoV-19/USA/NJ_3592/2020 | North America / USA / New Jersey | 2020-03-04 | NJ Public Health and Environmental Laboratories | Pathogen Discovery, Respiratory Viruses Branch, Division of Viral Diseases, Centers for Disease Control and Prevention | Yan Li, Krista Queen, Clinton R. Paden, Rachel Marine, Anna Uehara, Ying Tao, Jing Zhang, Haibin Wang, Mary S. Keckler, Alison S. Laufer Halpin, Christopher A. Elkins, Suxiang Tong |
| EPI_ISL_424901 | hCoV-19/USA/NJ_3201/2020 | North America / USA / New Jersey | 2020-03-04 | NJ Public Health and Environmental Laboratories | Pathogen Discovery, Respiratory Viruses Branch, Division of Viral Diseases, Centers for Disease Control and Prevention | Ying Tao, Clinton R. Paden, Jing Zhang, Krista Queen, Anna Uehara, Yan Li, Haibin Wang, Mary S. Keckler, Alison S. Laufer Halpin, Christopher A. Elkins, Suxiang Tong |

| | | | | | | |
|---|---|---|---|---|---|---|
| EPI_ISL_429968 | hCoV-19/France/HF1463/2020 | Europe / France / Hauts de France / Compiegne | 2020-02-21 | Centre Hospitalier Compiègne Laboratoire de Biologie | National Reference Center for Viruses of Respiratory Infections, Institut Pasteur, Paris | Mélanie Albert, Marion Barbet, Sylvie Behillil, Méline Bizard, Angela Brisebarre, Flora Donati, Fabiana Gambaro, Etienne Simon-Lorière, Vincent Enouf, Maud Vanpeene, Sylvie van der Werf, Raulin Olivia |
